# Statistical Considerations in the Design and Analysis of Longitudinal Microbiome Studies

**DOI:** 10.1101/448332

**Authors:** Justin D Silverman, Liat Shenhav, Eran Halperin, Sayan Mukherjee, Lawrence A David

## Abstract

Longitudinal studies of microbial communities have emphasized that host-associated microbiota are highly dynamic as well as underscoring the potential biomedical relevance of understanding these dynamics. Despite this increasing appreciation, statistical challenges in the design and analysis of longitudinal microbiome studies such as sequence counting, technical variation, signal aliasing, contamination, sparsity, missing data, and algorithmic scalability remain. In this review we discuss these challenges and highlight current progress in the field. Where possible, we try to provide guidelines for best practices as well as discuss how to tailor design and analysis to the hypothesis and ecosystem under study. Overall, this review is intended to serve as an introduction to longitudinal microbiome studies for both statisticians new to the microbiome field as well as biologists with little prior experience with longitudinal study design and analysis.

## Introduction

There is an increasing recognition that host-associated microbiota are a contributor to many aspects of human physiology and health and may ultimately play important roles in the diagnosis, treatment, and prevention of human disease (1-3). Beyond human health, the study of microbial community composition using high-throughput DNA sequencing have found use in a variety of fields including ecology, agriculture, and industrial engineering. The most common method for characterizing these communities is based on sequencing the 16S ribosomal RNA gene in bacteria which while ubiquitous in bacteria is also diverse enough to act as a molecular fingerprint distinguishing microbial taxa. The resultant data used for modeling and analyses of microbial community is a count table where each count reflects the relative abundance of a given taxa in a given sample. Considerable effort now focuses on best practices for designing microbial community surveys and methods for the analysis of such sequence count data.

An increasing appreciation of the temporal variability of host-associated microbial communities and the potential biomedical relevance of these dynamics has led to new statistical challenges (1, 4-8). For example, in Caporaso et al. (4), the authors study the natural variability of human microbiomes over the course of two years. Understanding such intrinsic temporal variability within microbial communities also requires statistical methods of partitioning observed temporal variation into technical and biological components (9-11). In contrast, in Yassur et al. (12), the authors focus on the temporal effects of antibiotics on infants gut microbiome in the first three years of life. Modeling such external factors requires dynamic models capable of including external covariates and potentially lagged or non-linear effects on the microbiota (13, 14). Alternatively, large longitudinal studies have been collected with the purpose of guiding diagnosis of inflammatory bowel disease and type 2 diabetes as well as prediction of preterm birth (15). Building such diagnostic tools will will require predictive models that can either forecast current or future community regimes based on the community composition at earlier time points are requires (10, 16).

The statistical considerations of longitudinal microbiome studies that could ultimately affect the biological findings, goes beyond data modeling and encompases study design, data preprosessing, and model evaluation. The data generation process underlying the profiling of microbial communities, using high-throughput sequencing *(e.g.,* sample processing and measurement), has implications for the design and analysis of the resulting data. We therefore begin this review of statistical considerations of longitudinal microbiome data with a thorough discussion of the measurement process underlying sequence counting (Section 1). Proper experimental design can be vital to the ultimate success of a study. As longitudinal studies impose an added layer of complexity beyond cross-sectional designs, we discuss the design of longitudinal microbiome studies in Section 2. Finally, we discuss the modeling of longitudinal microbiome data in Section 3. As data preprocessing and model evaluation are also crucial components in the analysis of longitudinal data, we discuss these steps as well as part of data modeling in Section 3. We highlight that our discussions regarding the data generation process and experimental design are largely novel with minimal consideration in previous reviews. This review is aimed towards applied statisticians unfamiliar with microbiome data and experimentalists with a minimal background in the analysis of longitudinal data.

## Sample Processing and Measurement

The majority of longitudinal microbiome studies to date have made use of high-throughput sequencing, where the relative abundance of different bacterial taxa is inferred based on the amount of distinct DNA reads. Yet, due to a wide variety of technical factors, such measurements of microbiota can differ substantially from the underlying true community structure (17). These discrepancies, which are not limited to longitudinal measurements, can be attributed to the sample processing-a sequential procedure, where in each step DNA is randomly selected, processed, and then included in the subsequent step. The randomness of this data generation process introduces uncertainty into microbiome measurements in the following way. First, this process results in a competition to be counted, or in other words more of one taxa results in less of another for purely technical reasons (Figure 2). Without knowledge of the total microbial load in the original ecosystem, this competition to be counted limits us to inferences regarding the relative abundance of taxa in that ecosystem (Figure 3). The second issue is that, while the read count data contains only relative information, the random sampling process also introduces uncertainty in estimates of these relative measurements that must be considered in analyses (18). Given the randomness in the data generation process described above, we suggest directly modeling the uncertainty in the counts using a model that also considers this competition to be counted. The multinomial distribution is the prototypical example of such a model. In particular, the multinomial distribution addresses count uncertainty rather than directly transforming counts to relative abundances and also models the competition to be counted rather than treating the counts of each taxa as independent. Additionally, the multinomial distribution may also better model the excess zero values often found in microbiome datasets that can be due to increase in the abundance of one taxa causing other taxa to fall below the detection limit of sequencing.

**Figure.1.**
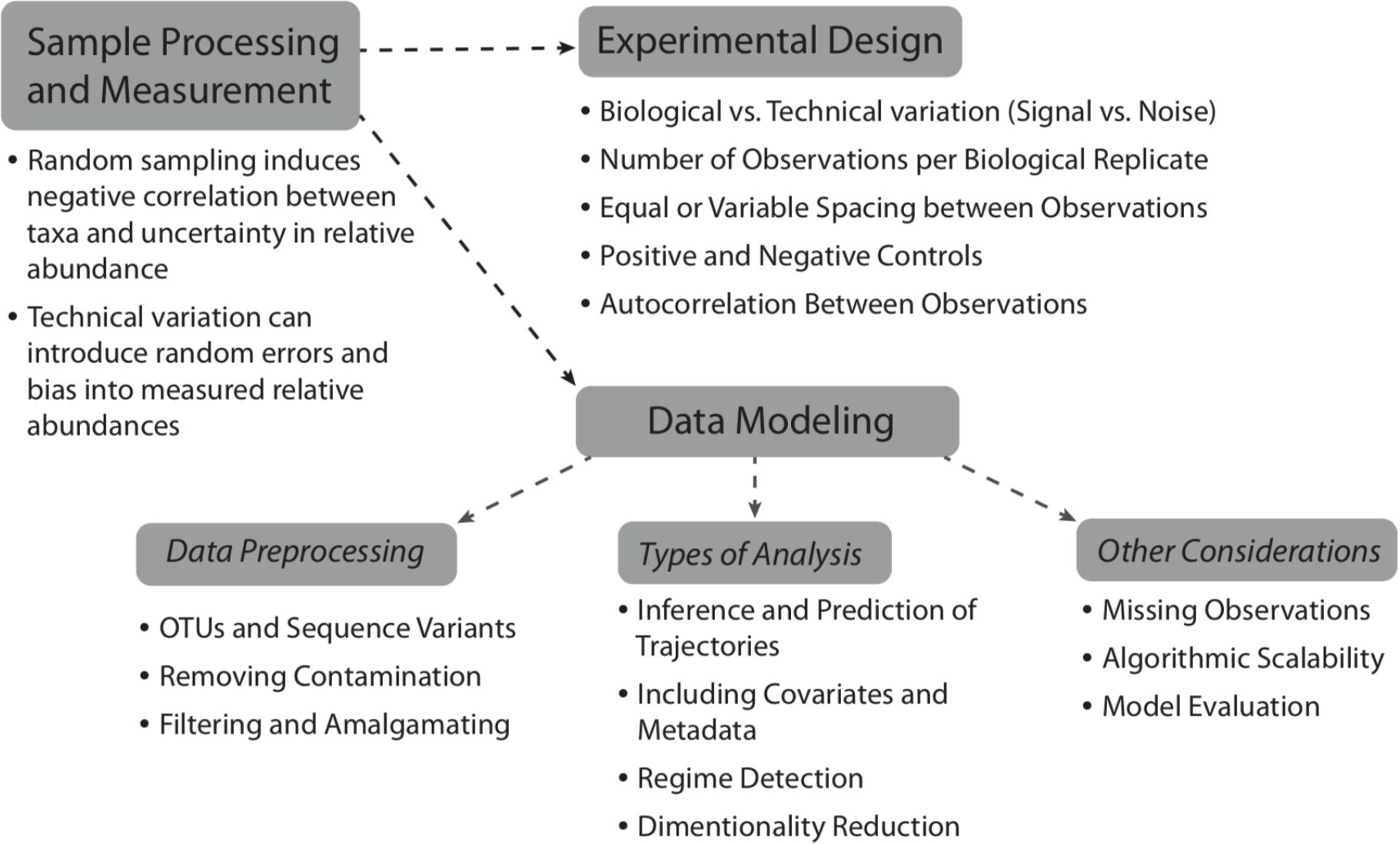
Structural Overview of this Review-statistical considerations of longitudinal microbiome data with a thorough discussion of: the measurement process underlying sequence counting (Section 1), the experimental design of longitudinal microbiome studies (Section 2) and modeling of longitudinal microbiome data (Section 3).

**Figure.2.**
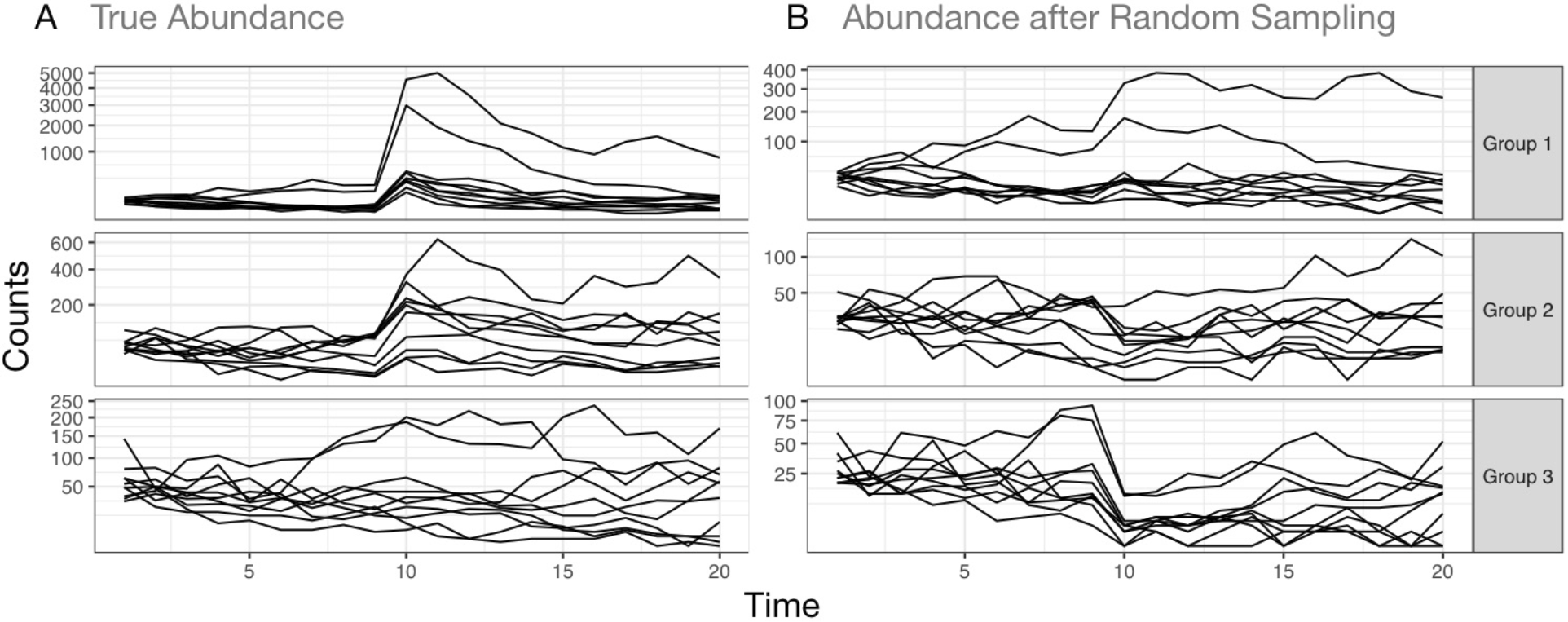
Effects of random sampling can lead to spurious conclusions in longitudinal studies. (A) A longitudinal study of 30 microbial taxa was simulated over 20 time-points. At time 10 a simulated prebiotic was given. 10 taxa in group 1 were simulated to grow rapidly in response to the prebiotic. 10 taxa in group 2 were simulated to grow moderately (half the response of group 1) in response. 10 Taxa in group 3 were simulated to have no response. (B) Each sample was randomly sampled to even sequencing depth and the resulting counts. This type of subsampling simulates the sample pooling step performed in multiplex sequencing where the DNA from each sample is subsampled to even depth to provide even coverage of sequencing depth across samples. Notably, bacteria in group 2 and 3 now display a marked and spurious decline in response to the prebiotic. Even the majority of those bacteria in group 1 that did not have the largest effect now appear to show no increase in response to the prebiotic. This effect reflects the competition to be counted that is introduced by random sampling.

**Figure.3.**
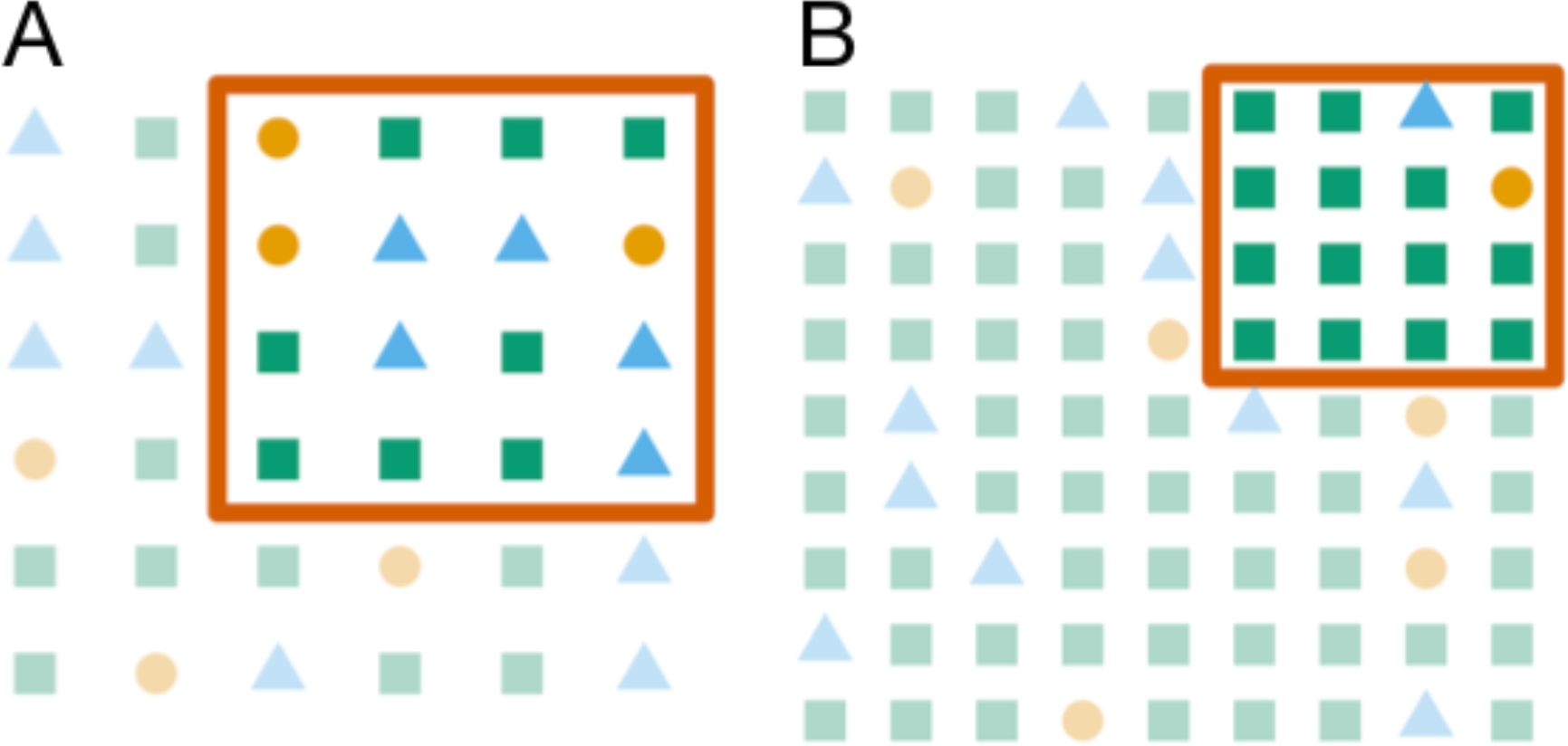
Random sampling limits inferences to the relative abundances of different microbial taxa and induces negative correlation into the observed data that may lead to spurious conclusions. Panels A and B illustrates sampling (red square) from two microbial ecosystems with three types of bacteria (depicted as orange circles, green squares, and blue triangles). In Panel A, measuring three orange taxa reflects only an arbitrary sample size (not the absolute abundance of the orange taxa). In contrast, measuring three orange and five blue taxa provides information on the ratio of orange to blue taxa in the entire ecosystem. Panel B depicts sampling from an ecosystem that is identical to that in Panel A but with an increase in the absolute number of the green taxa. As overall sampling size is fixed, it is likely that a random sample will count fewer orange and blue taxa and more green taxa. Compared to Panel A researchers may then be led to believe that Panel B contains more green and less blue and orange taxa whereas in reality only the green has changed. Even if the sample size in Panel B was not the same as in Panel A spurious conclusions may be reached.

Notably, while the multinomial distribution can account for the uncertainty stemming from random sampling, it alone is not sufficient to model other sources of technical variation in the observed data. Specifically, studies have found that numerous steps including DNA extraction, PCR amplification, DNA sequencing, sample collection and even sample storage can impact measured microbial composition (17, 19-24). To capture these forms of technical variation, and to model other biological sources of variation, we recommend models that also account for these other sources of variability either through extensions of the count distribution, such as the Generalized Multinomial-Dirichlet, or as we describe below by modeling variation in the parameters of the count distribution.

While our discussion to this point has focused on approaches to accurately model microbial relative abundance from sequence count data, recent methods have proposed approaches that aim to produce measurements based on microbial absolute abundances using external measurements (25-27). Two methods in particular, qPCR and Flow cytometry have gained increasing popularity for this purpose (25). Yet, accurately recovering the absolute abundance of taxa in an ecosystem requires direct measurement of total microbial load which is often not possible *in vivo.* Instead, available methods provide an estimation of the microbial load at some later sample processing step that does not necessarily reflect the ecosystem total but rather the microbial concentration in the stool. Nonetheless, use of such external measurements to augment relative abundances inferred from sequence count data should still consider modeling count uncertainty and competition to be counted as well as measurement error present in the extraneous total information.

## Experimental Design

Experimental design can be key to the success of any study. Notably, longitudinal studies require additional parameters be chosen beyond those required for standard crosssectional designs. In particular, the parameters involved in the design of longitudinal microbiome studies include the number of individual time-series to study (e.g., the number of individuals studied) !, the number of time-points in each time-series, T_r_ (for r = 1, …,R), the number of taxa), and the number of measured covariates * *(e.g.,* diet, sex, antibiotics, or batch number). The design of longitudinal studies must also consider the spacing (either equal or variable) between time-points within each time-series and the synchronization or lack thereof of samples between individuals. Additional measurements in the form of technical replicates or positive and negative controls may also be required to adequately partition biological signal from technical noise. As a final complication, the design must be motivated and account for subtleties of the ecosystem under study and the research goals. However, ultimately the general principle underlying all experimental design is the need to differentiate biological variation *(i.e.,* signal) from technical variation *(i.e., noise).* For this purpose, we recommend thinking about the interplay between technical variation, biological variation, and the temporal distribution of sample collection.

As sampling effort of biological systems are often limited both by cost and by biological limitations (e.g., the rate at which an organism defecates), care is needed to appropriately distribute sampling effort. The primary goal when portioning sampling effort is to ensure that sampling frequency is adequate to capture the biological variation of interest (7, 28). One way to address resource limitations is to concentrate sampling effort around highly variable time-periods such as right before and after an experimental intervention or using prior knowledge on the ecosystem’s dynamics. Yet, modeling such irregularly spaced studies can be challenging, requiring special modeling techniques such as state space models or imputation methods. A guiding principle when designing irregularly spaced studies is to aim to achieve equal amounts of variation between consecutive time-points (29). Additionally, if it is known that the biological variation of interest is periodic and no aliasing is occurring then autocorrelation can be a useful tool for determining sampling rate and sampling spacing. In this situation, one should consider sampling at twice the frequency of the signal (30).

Given that sampling effort is portioned appropriately to capture the biological variation of interest, the impact of technical variation must be considered. In particular, it can be helpful to consider two forms of technical variation, random errors and systematic biases (31). Random errors, such as pipetting errors, are those sources of variation that cause biologically identical samples processed in parallel *(i.e.,* technical replicates) to differ. Systematic biases, such as a consistent underrepresentation of Actinobacteria in measured data (32), are those sources of variation that cause technical replicates to differ from the truth but remain nearly identical to each other. If the relationship between technical and biological variation is unknown or technical variation is known to be at least comparable to biological variation, special care is needed to account for technical variation.

For random errors, the two standard solutions are to either collect ample technical replicates or to sample more frequently. While the former always works the later can encounter similar biological or cost limitations on sampling frequency and will also require an assumption that longitudinal trajectories varying smoothly and slowly compared to the sampling frequency. For example, Silverman et al. (9) quantified and control for random errors (and not systematic biases) in an in vitro system by using ample technical replicates which allowed the characterization of the relationship between technical and biological variation as a function of sampling frequency for an in vitro artificial gut system. They found that at hourly sampling frequencies technical variation was approximately four times larger than biological variation, and was equal at sampling frequencies of approximately 3.5 hours. These results demonstrated that subsequent studies in these ecosystems must either account for technical variation or choose to limit inference to biological variation occurring over longer time-periods (e.g., daily).

As for systematic biases, few solutions are available as measuring and correcting these biases requires either specially designed calibration experiments or positive and negative controls. A notable example is the work of Davis et al. (33) who proposed a method for removing contamination in microbiome sequencing studies using designed negative controls. Nonetheless, capturing other sources of technical bias such as PCR bias or DNA extraction bias remain outstanding problems. On the positive side, systematic biases, unlike random errors, may be ignored in many situations where the specific aims of the study are not affected by such technical variation. For example, if the aim of a study is to identify how the community structure changes over time, a systematic bias such as the consistent underrepresentation of Actinobacteria in observed data, may not affect inference. In contrast, if the aim of a study is to determine which taxa is the most abundant at any given time-point, such underrepresentation may severely effect inference.

Often a chief concern in data analysis is that technical variation covaries with a biological feature of interest (34). Specifically, in longitudinal studies sample randomization throughout sample processing can mitigate issues that may arise from covariation between technical variation and the sample time-index. Additionally, all sample processing information should be retained and reported with the final dataset as various modeling approaches may be used to correct for differences between groups of samples processed separately such as linear mixed models (35).

As the problem of experimental design for longitudinal microbiome studies is challenging, model based methods for optimizing the study design may provide substantial insights. In particular some of the most basic questions such as how to choose values for R and *T* using sample size or power calculations remain under-studied. Of the few available techniques, we recommend using simulation studies. As in power and sample size calculations, simulation studies can provide a simple (yet computationally intensive) method for answering such questions. For example, Fukuyama et al. (36) used simulation studies to conclude that for their research goals, crossover longitudinal sampling designs with baseline correction are more powerful than parallel designs. In the same paper the authors, designed their irregular sampling intervals to achieve approximately equal variance between time-points using simulations studies. Despite the utility of simulation studies, formal methods of experimental designs such as those maximizing D-optimality criteria, information gain, or those based on Bayes optimality may provide methods of optimizing many more aspects of experimental design (37). To the best of our knowledge, the only available methods for optimal design of longitudinal microbiome experiments are those provided in the MC-TIMME package (13) which selects a sampling frequency to maximize the expected information gain.

Finally, additional considerations in the design of longitudinal microbiome studies, such as the temporal relationships between time-series or with respect to external events, should be appraised. Studies involving multiple individuals over time (R > 1) should design the experiment in a way that will subsequently be utilized by the the data modeling scheme. For example, multi-person studies of the effect of a targeted intervention may want to collect samples at similar temporal distances from the intervention. Such synchronization allows for richer questions to be investigated and greatly reduces modeling complexity. Seasonality effects can also be important to consider in experimental design. While perhaps most obvious in environmental studies, dietary changes surrounding holidays or weekends, or even natural circadian variation (38) can also be important factors to consider.

## Data Modeling

Due to the complexity and high-dimensional nature of longitudinal microbiome data, nearly all datasets will require non-trivial statistical modeling. In this section we discuss various aspects of data modeling including the initial preprocessing of data, various types of analyses that are commonly performed, and various considerations in choosing models.

### Data Preprocessing

Two approaches are commonly used to characterize microbial communities by summarising raw sequencing reads into a sequence count table: quality filtering OTU-based methods and denoising methods. In the OTU-based methods, all the sequences are clustered into OTUs based on a distance matrix at a specified dissimilarity threshold (typically 3%), which reduces the rate at which errors are misinterpreted as biological variation. However, OTUs underutilize the quality of modern sequencing by precluding the possibility of resolving fine-scale variation. The denoising methods, e.g., DADA2, model the error process and evaluate the validity of individual sequences in the context of the full metagenomic data set, while including the number of reads corresponding to each sequence. Notably, a growing consensus has arisen that methods like DADA2 help avoid the inflation of diversity often seen with many OTU clustering methods (39). Using DADA2 improved resolution of low-frequency taxa may be achieved by pooling or pseudo-pooling which uses a two pass algorithm to adaptively threshold low-abundant sequences by pooling information across samples from similar environments.

Despite the significant progress in reducing errors from PCR and sequencing, the accuracy of microbial community surveys still suffers from the presence of contaminants — DNA sequences not truly present in the sample. Preparing contaminant-free DNA is challenging, and the sensitivity of PCR and whole-genome amplification methods means that even trace contamination can become a serious issue (40, 41). To alleviate this issue, computational methods for ‘microbial source tracking’ (quantifying the contribution of potential source environments to complex microbial communities) have been proposed (33, 42, 43).

These methods identify both the source and quantity of contamination, and could help account and remove the contamination. Another option to account for contamination is using taxonomy-dependent methods. These methods rely on the annotated sequences already deposited in the databases for taxonomic assignment of a query sequence by the best-matching sequence in the reference database. Although taxonomy-dependent methods can assign taxonomy to the query sequences based on previously characterized microbes, lack of sufficient well-characterized microbes and reliable taxonomy often make it difficult to characterize novel sequences, and the robustness and accuracy of such methods are mainly dependent on the completeness of the annotated reference database. The count table that results from either OTU clustering or denoising methods is typically very sparse, often with greater than 90% zero values, which can pose both computational and statistical problems (44). In particular, estimates of relative abundance from small counts involves large variance and therefore low certainty (45). To focus analysis on taxa whose relative abundance can be estimated with higher certainty and to alleviate the computational burden of excessive sparse taxa, analyses typically filter low abundance taxa prior to modeling. However, the thresholds used for filtering are typically heuristics and have not been thoroughly investigated. In determining whether initial data filtering is appropriate, and if so how it will be done, the goals of a study must be considered. In general, studies aimed at exploratory analyses, such as biomarker discovery, should consider little to no initial filtering. In particular, such studies may consider filtering taxa based on their temporal patterns as demonstrated by Shenhav et al. using the time-explainability measurement they introduced, which corresponds to the fraction of variance explained by the microbiome composition in previous time points. Specifically, time-explainability is informative for selecting time-dependent taxa (i.e., taxa that can be predicted based on the previous microbial composition). In contrast, for studies aimed at confirmatory analysis we suggest limiting the number of taxa analyzed to just those groups under study. For example, if the purpose of a study is to determine the dynamics of butyrate producing organisms, researchers should consider amalgamating taxa into butyrate producing and butyrate non-producing groups and restricting analysis to just these two categories. Overall, as the total number of counts in each sample provides important uncertainty information, we recommend that studies never eliminate counts/taxa from a dataset but instead amalgamate taxa that are not of primary interest into a category called “other”.

### Types of Analysis

In this subsection we focus on modeling the temporal relationship between samples towards goals such as inference and prediction of microbiota longitudinal trajectories. A common task in the analysis of longitudinal microbiome data is to infer the trajectories of the community as well as forecasting future compositions. Longitudinal microbiota data may include many different temporal patterns such as cyclical effects *(e.g.,* seasonal or circadian effects), long term trends, or even delayed effects of shifts in community composition. However, models may differ substantially in the types of temporal patterns they can infer and predict. For example, the model of (14) is capable of capturing single time-step interactions. Other models such as the Poisson ARIMA model of Ridenhour et al. (46) allow for temporal interactions to be carried over multiple time-steps. Even greater flexibility is allowed by models such as those of Shenhav et al. (10) and Silverman et al. (9) which allow for more complex time-series modeling such as the inclusion of seasonal or polynomial trends to be included as well. Other methods can achieve even greater flexibility by using non-parametric kernel methods to either model community trajectories using Gaussian processes (11) or finding low dimensional representations of trajectories (36) (although the later does not, strictly-speaking, enable prediction or account for the temporal correlation between samples). However, these nonparametric methods may not allow for parameter estimation of distinct temporal components as parametric methods can *(e.g.,* to quantify the relative impact of seasonality versus long term trend).

Another common goal is to infer or predict the relationship between external metadata (*e.g.,* covariates or perturbations) and microbiota trajectories, including the effect of perturbations. For example, the differences in temporal trajectories between treatment and control groups in a longitudinal study may be investigated by incorporation of a binary covariate indicating presence of a measurement in either a treatment versus control group. Alternatively, inclusion of time-varying metadata such as the pH of the environment can be used to explore the impact of such dynamic factors on a microbial community. The methods of (9, 10, 14) all allow for such external covariates. In particular the MALLARD class of models introduced by Silverman et al. (9) further allow for the effects of such external covariates to be time-varying as well, which is a modeling procedure known as dynamic regression (47). While these methods allow for very flexible modeling of linear interactions between covariates and microbiota trajectories, non-linear modeling of perturbations as is required popular in classical pharmacokinetic (PK) and pharmacodynamic (PD) studies remains understudied. Multivariate extensions of standard PK-PD models or non-linear transfer function models would likely find tremendous use in studies aiming to understand the impact of therapeutics on the microbiome.

Of course the above is just a limited sampling of common analysis tasks. Other tasks include regime or changepoint detection, dimensionality reduction or feature selection, and time-series synchronization. First, regime and changepoint detection relates to the problem of either identifying a regions within a time-series that differs in some substantial way from other parts of the time-series. While few methods have been adapted to address the many specific challenges of microbiome data, Sankaran and Holmes (16) provide a thorough review of available methods and recommendations on productive new research directions. Second, as longitudinal microbiome data is high dimensional, with studies typically analyzing tens to thousands of taxa, methods for dimensionality reduction or feature selection can greatly improve scalability and interpretability of models. A small number of methods to date have addressed these issues and include the post-processing dimensionality reduction based low dimensional representations of biological temporal covariation inferred by state-space models and introduced in Silverman et al. (9); dimensionality reduction using Kernel ordination as in (36); Dirichlet process clustering of temporal patterns introduced in Gibson and Gerber (14); as well as autoregressive-based feature selection using linear mixed models introduced by Shenhav et al. (10) which aims to retain taxa whose trajectories are most explainable by time. Finally, time-series synchronization refers to the problem of aligning temporal patterns in distinct time-series or aligning time-series from unsynchronized experimental designs. Specifically, temporal alignment for microbiome data was suggested by Lugo-Martinez et al. (48) as a preprocessing step, that may improve the prediction accuracy of a Dynamic Bayesian network, similar to the model suggested by Larsen et al. (49).

### Other Considerations in Model Choice

Missing observations in longitudinal microbiome studies are common and can pose a challenging modeling task by essentially interrupting the temporal chain of observations. Observations missing due to extraneous factors, such as issues arising during sample processing, are often termed missing at random and may be handled in a number of ways. Perhaps the most common means is by concatenating together samples on either side of missing observations. This practice however is discouraged as it can lead to biased inference, often increasing the inferred temporal variation especially for studies with multiple or consecutive missing observations. Instead we recommend that observations missing at random be modeled directly either through marginalization as was performed by Silverman et al. (9), or through the use of non-parametric kernel methods that concern themselves only with the distance (or conversely similarity) between observations as is the case with the methods of Aijo et al. (11) and Fukuyama et al. (36). In contrast to observations missing at random, observations may be missing for other reasons that may confound inference, such as censoring due to subject withdrawal from a study due to adverse drug reactions. This later type of missing observations is often termed missing not at random and often requires a combination of expert knowledge and special modeling techniques that are beyond the scope of this review.

Scalability is another important factor of successful inference and prediction in longitudinal microbiome studies. Importantly there is a balance between the scalability of a model and the assumptions made, where, the more covariation between parameters in a model and the more levels of latent variables, the more computationally intensive to infer or predict. In Table 1 we highlight assumptions made by each model. The difference in scalability between existing models can be large with only a subset of models providing results in a reasonable time-frame. However, such assumptions must be considered carefully as subtleties of these assumptions can have large impacts on results. Specially, only the models proposed by Shenhav et al. (10), Ridenhour et al. (46) and Fukuyama et al. (36) can be applied on the entire microbial community, at any level of the phylogenetic tree (taxa/genus/order/family etc), while all other methods mentioned above require either some feature selection in the taxa level (<200) or amalgamation to a higher phylogenetic level (genus/family/order).

**Table 1.** Comparison of six currently available tools for modeling longitudinal microbiome studies. Other notes: 1) adaptive gPCA does not provide a method of prediction, but rather a low-rank representation; 2) only GP Microbiome and MALLARD model a competition to be counted in the sampling process; 3) GP Microbiome can only model stationary dynamics as a result of its radial basis function kernel.

| Name and Summary | Tasks | Measurement Model | Highlighted Assumptions | Software | Scalability | Missing Data | Multiple Time-Series |
| --- | --- | --- | --- | --- | --- | --- | --- |
| **Poisson ARIMA** (46) | Inference and Prediction of long-term trends, high-order temporal dependency | Poisson | (1) No relative constraints<br>(2) Dynamics are Log-Linear | ARIMA(1,0,0)<br>(only) offered by the authors upon request | ++ | No | No |
| **MTV-LMM** (10) - Linear Mixed Models with Variance Components | Inference and Prediction of long-term trends, high-order temporal dependency, and metadata covariates; Feature selection | Transformed Multivariate Normal (e.g., Standard, Log, or Logistic) | (1) No count noise (2) Dynamics are Linear or Log-Linear | R Package | ++ | Yes | Yes |
| **Adaptive gPCA** (36) | Inference of Low dimensional structure in observed data | Transformed Multivariate Normal (e.g., Standard, Log, or Logistic) | (1) No count noise (2) No temporal correlation between samples (3) User defined kernel | R Package | ++ | Yes | Yes |
| **MDSINE 2.0** (14) - Bayesian Ecological Interaction Model with Clustering | Inference and Prediction of First-order autoregressive trajectories with metadata covariates; Clustering taxa based on dynamics | Negative Binomial for counts, Univariate Normal for qPCR | (1) qPCR reflects microbial load (2) First order temporal dynamics only (3) Dynamics are Log-Linear | None | + | Yes | Yes |
| **GP Microbiome** (11) - Bayesian Generalized Gaussian Processes | Inference and Prediction of any smoothly varying trajectory | Multinomial - Logistic Normal | (1) Zero Inflation "Shot Noise" is present and independent for each count in the dataset. (2) Stationarity | Command Line interface to Stan | - | Yes | No |
| **MALLARD** (9) - Bayesian Generalized Dynamic Linear Model | Inference and prediction of long term trends, high-order temporal dependency, metadata covariates, and identification and denoising of technical variation and bias | Multinomial - Logistic Normal | (1) Dynamics are Log-Linear | Stan code implementing MALLARD with time-invariant covariance | - | Yes | Yes |

Model evaluation is an important step in developing models as well as assessing their utility either as inferential or predictive tools. As ground truth is often unknown in many studies, statistical methods based on either model predictive accuracy or goodness of fit to observed data are often employed. With regards to longitudinal microbiome studies predictive metrics such as cross-validation which iteratively fits a model to a subset of the data and then assess the predictive accuracy of the fit model on the held-out samples are particularly useful. Potential metrics to assess the predictive accuracy of a model include the correlation between estimated relative abundance and the relativised counts or the distance *(e.g.,* Euclidean, Aitchison, or arcsine) between predicted relative abundance and relativised counts (10, 11). The first measurement highlights the model’s ability to accurately predict the abundance along time per taxa, while the second measurement highlights the model’s ability to accurately predict the microbial community composition per time point. Alternatively, statistical quantities such as the marginal log-likelihood, AIC, BIC, or WAIC can be used to assess the goodness of fit of a model to observed data (50). Typically such quantities include a penalization based on model complexity to mitigate overfitting.

## Conclusions

Here we have presented a review of statistical considerations in the experimental design and analysis of longitudinal microbiome studies. In addition to describing aspects of design and analysis we also presented a thorough discussion of challenges arising from the measurement process underlying sequence counting. In particular, we introduced how the process of sampling can introduce a negative correlation between counts *(i.e.,* competition to be counted) into observed data and a consideration of technical variation and bias that can also be present. As part of a larger discussion on design of longitudinal microbiome studies, we introduced concepts relating to the distribution of sampling effort along time as well as the interplay between biological and technical replicates. Finally, as part of our discussion of data modeling we discussed considerations in data preprocessing, analytic goals, and other considerations such as missing data, scalability, and model evaluation.

Although we have attempted to provide general guidelines regarding the design and analysis of longitudinal microbiome studies, these decisions must ultimately be based on the hypothesis and goals of a given study. For example, consider the following two studies. Study A aims to identify specific microbes in a microbial community that change cyclically along host day-night cycling (38). In contrast, study B aims to identify compositional shifts of microbial genera in patients undergoing allogeneic hematopoietic stem cell transplantation with concurrent antibiotic prophylaxis (51). Differences in the goals in studies A and B should lead to differences in how they approach technical noise, sampling frequency, data preprocessing, and modeling. For example, while study A is concerned with sub-daily variation and should likely sample at least every 6 hours, study B is more concerned with longer term trends and likely only needs to sampled daily. Additionally, we would expect that the effects of antibiotic treatment in study B would lead to larger compositional shifts than the natural day-night cycling in study A. Therefore, study A should be more concerned with the impact of technical variation than study B. Finally, whereas study A aims to identify, with highest resolution, the taxa that may oscillate with host day-night cycles, study B is specifically interested in shifts at the level of bacterial genera. Therefore whereas study A should consider performing inference at the level of sequence variants or OTUs, study B should likely amalgamate taxa to the level of bacterial genera to improve statistical power and decrease computational complexity.

Challenges in the design and analysis of longitudinal studies introduce many new avenues for future research. Perhaps the most pressing need is for scalable models that account for count variation and the negative correlation between taxa introduced by random sampling. While models based around the multinomial distribution account for these features, existing implementations are not scalable to larger microbiome analyses. Second, the majority of existing methods have focused on trajectory inference or prediction. Methods that address other questions such as trajectory classification would be of great value. Finally, there is a need for models that account for delayed or multi-time-step effects of perturbations as occur in pharmacokinetic or pharmacodynamic studies. Few current models can easily handle such perturbations and as such multivariate PK-PD models or models that incorporate non-linear transfer functions would fill a current void.

